# MetaDome 2027: a comprehensively updated resource for aggregating missense variant evidence across homologous human protein domains

**DOI:** 10.64898/2026.08.26.747388

**Authors:** Laurens Wiel, Federico Ferraro, Jay Yu, Jimmy Zhen, Daniel Nachun, Rodrigo Mendez, Chloe M. Reuter, Jason L. Cui, Devon E. Bonner, Jennefer N. Carter, Shruti Marwaha, Maartje van de Vorst, Sara Emami, Elijah Kravets, Matthew B. Neu, Tjakko W. van Ham, Tjitske Kleefstra, Euan A. Ashley, Jonathan A. Bernstein, Stephen B. Montgomery, Christian Gilissen, Matthew T. Wheeler

## Abstract

The interpretation of missense variants remains a major challenge in clinical genetics. “Meta-domains” aggregate population and pathogenic variation across homologous Pfam domain instances in the human proteome, providing per-residue context for interpreting variants of uncertain significance (VUS). Our 2019 implementation, MetaDome, is widely used and named in clinical variant-classification guidelines. Here we present the MetaDome 2027 update, featuring a comprehensively updated dataset and GRCh38 support. The redesigned pipeline enables incremental updates of GENCODE, UniProtKB/Swiss-Prot, Pfam, gnomAD, and ClinVar while maintaining 100% sequence-identity gene-to-protein mapping. Annotated Pfam domain instances grew 14.9% from 71,419 to 82,069 and meta-domain-eligible Pfam families (≥2 human occurrences) by 73.3% from 3,334 to 5,778; Pfam domains are annotated to 92% of human proteins. Approximately 43% of mapped protein-coding nucleotides (14.3 million in GRCh38, 13.8 million in GRCh37) are in a meta-domain; in GRCh38 67.9% (37,692 of 55,548) of pathogenic or likely pathogenic ClinVar missense variants fall at such a position. We show how MetaDome helped reclassify a *de novo* missense VUS in *RALA* and identify 52,463 ClinVar missense VUS for which meta-domains supply otherwise unavailable pathogenic evidence. MetaDome is freely available at www.metadome.app.

**Graphical Abstract:** 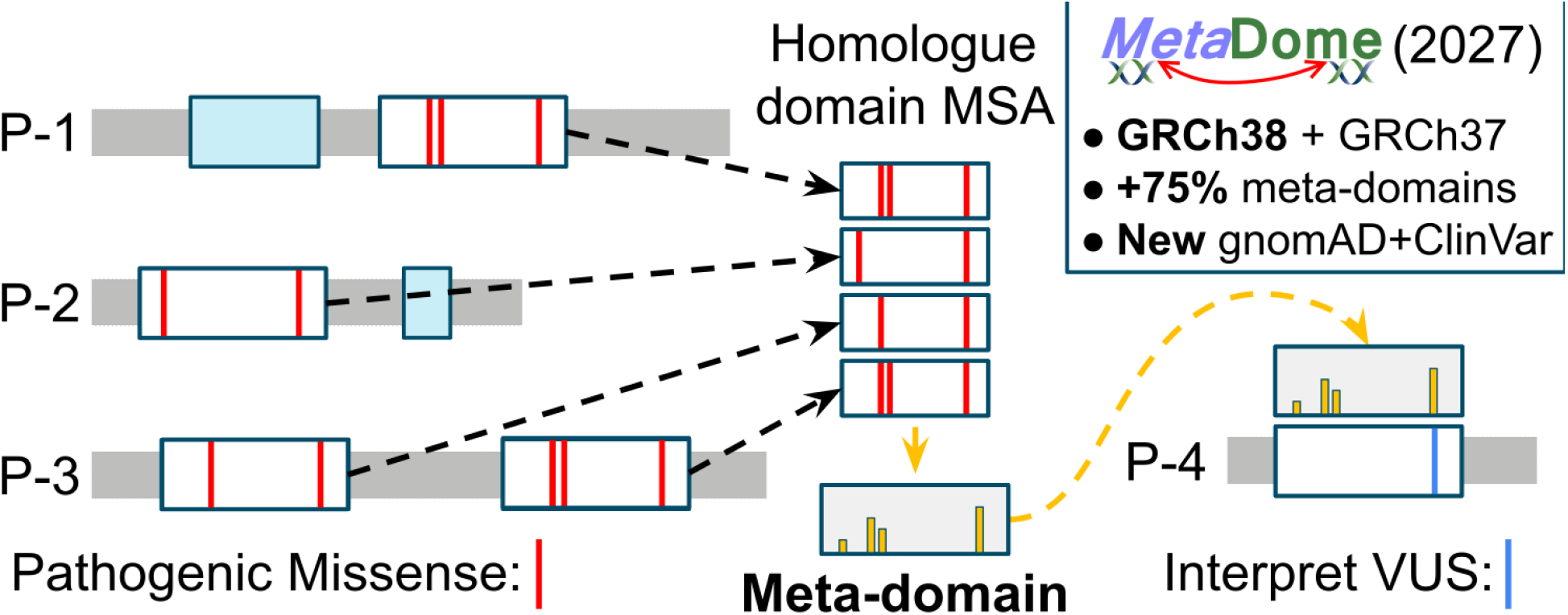

## Introduction

The continuous accumulation of human genomic data has driven the development of methods that interpret genetic variants by their context in large reference datasets. Population-frequency catalogues like gnomAD (1), pathogenic-variant databases such as ClinVar (2) and HGMD (3), gene- or region-level intolerance metrics such as RVIS (4), subRVIS (5), the Missense Tolerance Ratio (6), the missense observed/expected ratio (7), and our MetaDome Tolerance Landscape (8), are now standard inputs into clinical variant interpretation. Deep-learning predictors trained on evolutionary and structural information, like AlphaMissense (9), EVE (10), and PrimateAI-3D (11), have further improved per-variant scoring. However, most observed missense variants are individually rare, and a single observation provides little information (1).

One way to amplify the per-position signal is to look at homologous positions across proteins. Mutations at corresponding locations in homologous proteins tend to have similar effects on protein stability and function (12), and disease-relevant information transfers between paralogous disease genes (13). Building on this principle, we previously aggregated population and pathogenic variation across homologous Pfam protein domain instances within the human proteome and showed that pathogenic missense variants at equivalent positions in such “meta-domains” are often paired with the absence of population-based variation, and vice versa (14). We then exposed this concept for a wider range of adoption through MetaDome (8). The methodology was subsequently extended in Meta-Domain HotSpot (MDHS) (15), which used proteome-wide meta-domain based aggregation of *de novo* mutations from a large cohort of individuals with neurodevelopmental disorders to identify function-altering missense hotspots. Independently, the concept was formalised through a Bayesian variant-classification framework showing that pathogenic missense variants at an equivalent meta-position in a different protein provide moderate evidence of pathogenicity, with corresponding moderate evidence of benignity for benign meta-position variants (16). Meta-domains were also shown to improve prioritisation of *de novo* missense variants in developmental-disorder cohorts through a Homologous Missense Constraint (17). Together these independent studies establish the meta-domain principle as an evidence type for missense variant interpretation. That evidence has since been incorporated into the operational guidance now followed by clinical diagnostic laboratories (see Community Use of MetaDome).

Since the 2019 release of MetaDome, the underlying public datasets have grown; gnomAD has expanded from ∼123,000 exomes to over 800,000 combined exomes and genomes, and pathogenic and likely pathogenic missense variants in ClinVar considered by MetaDome grew from ∼47,300 to over 55,500. GENCODE has annotated thousands of additional protein-coding transcripts. Pfam has added new domain families and refined existing ones (18). The reference assembly has shifted: clinical sequencing pipelines now overwhelmingly call against GRCh38, while a substantial body of historical clinical sequencing data is still anchored to GRCh37. The original MetaDome implementation predated all of these changes.

Here we present a comprehensive update of the MetaDome platform and its underlying mappings. We rebuilt the mapping pipeline, integrated the latest releases of GENCODE, UniProtKB/Swiss-Prot, Pfam, gnomAD, and ClinVar, added support for GRCh38 alongside GRCh37, exposed a positional URL endpoint (**Figure 1A**) that supports direct query from external pipelines, and modernised the visualisation layer (**Figure 1B**). We then illustrate what the updated resource makes possible: a *de novo* missense variant in *RALA*, reclassified from uncertain significance to likely pathogenic in a diagnostic laboratory with support from pathogenic variants at the equivalent meta-domain position in *RIT1* and *KRAS*; and, proteome-wide, 52,463 ClinVar missense variants of uncertain significance that sit at meta-domain positions carrying uncontradicted pathogenic evidence and have no pathogenic variant at the residue itself. The per-residue dataset carrying tolerance scores and meta-domain aggregated variation, the underlying transcript-to-protein-to-domain mapping database, genome-browser tracks for both assemblies, and the prebuilt output needed to reproduce the MetaDome platform’s analyses are deposited publicly on Zenodo as standalone, reusable resources.

**Figure 1.**
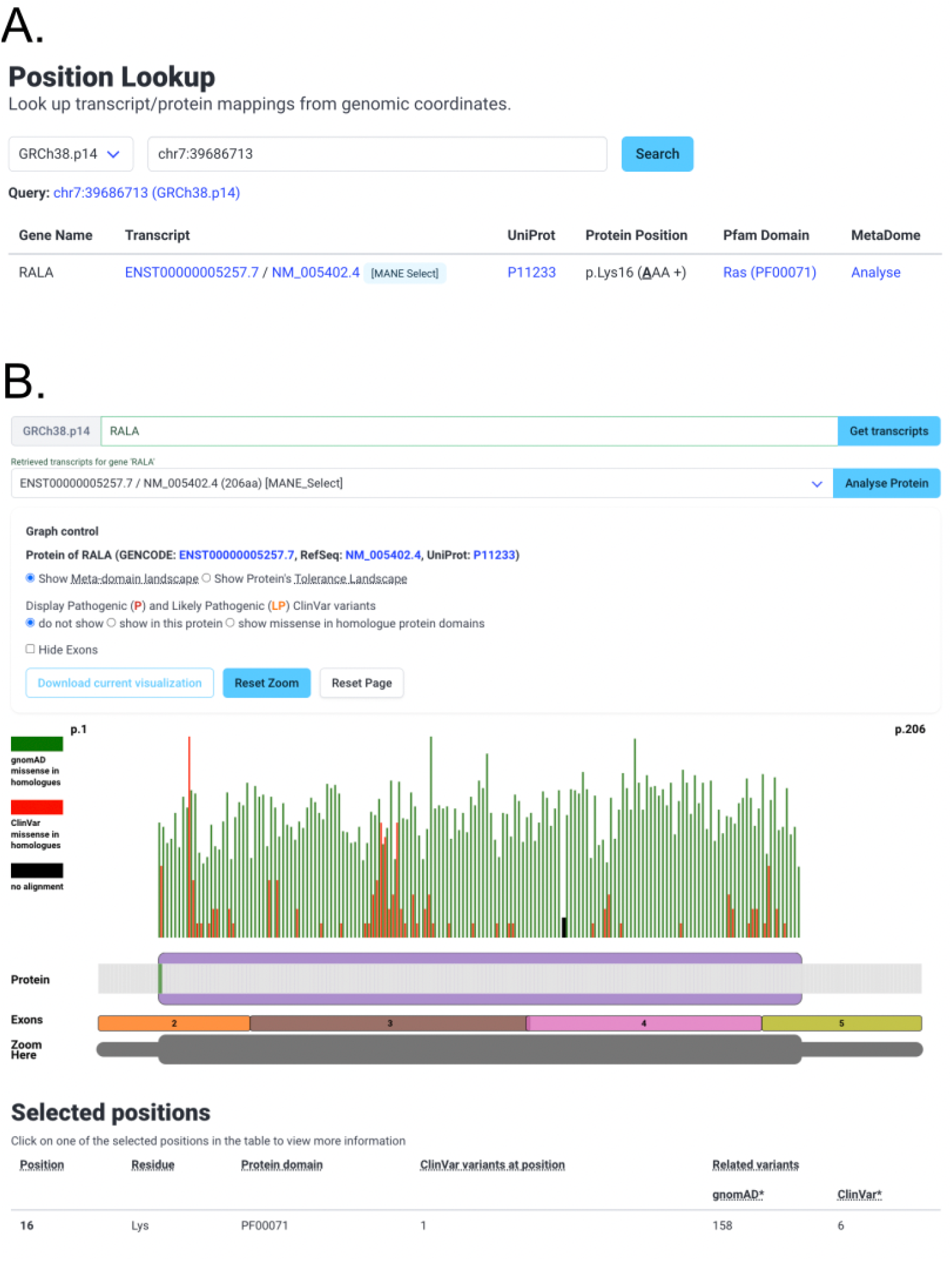
The MetaDome 2027 web interface. (**A**) The position-lookup page, new in this release, resolves a genomic coordinate to every transcript and protein context covering it. The user selects a reference assembly and enters a coordinate; each result row gives the gene symbol, the GENCODE and RefSeq transcripts with MANE Select status where applicable, the UniProt accession, the protein position with its codon and strand, any covering Pfam domain, and a direct link into the dashboard at that residue. (**B**) The per-protein dashboard, shown for RALA (ENST00000005257.7, 206 residues) with p.Lys16 selected. The layout follows MetaDome 2019: a gene and transcript selector, a graph control panel, the meta-domain landscape, a per-position selectable schematic protein view with Pfam domains, a zoom control, and a table of selected positions. New in the 2027 release are the reference-assembly selector, which allows GRCh37 and GRCh38 to be queried from the same interface; MANE Select annotation of transcripts; an exon track beneath the protein schematic with a control to hide it; residue-position labels bounding the landscape; and a ClinVar display control that is explicit about Pathogenic and Likely Pathogenic classification and offers variants in this protein and variants in homologous domains as separate choices rather than independent checkboxes. For p.Lys16 the selected-positions table reports one ClinVar record at the position itself and six at homologous meta-domain positions, the evidence described in the illustrative use case.

### Database Content and Data Sources

MetaDome integrates five primary public data sources: GENCODE (19) for protein-coding gene structure and translation sequences, UniProtKB/Swiss-Prot (20) for canonical and isoform protein sequences, Pfam (18, 21) for protein-domain Hidden Markov Models (HMMs), gnomAD r2.0.2 (7) and v4.1 (1) for population variation, and ClinVar (2) for clinically curated pathogenic variation. Reference assembly support is provided in parallel for GRCh37 (GENCODE v19) and GRCh38 (GENCODE v45). UniProtKB/Swiss-Prot release 2025_01, Pfam v37.4, gnomAD v4.1 (GRCh38) and r2.0.2 (GRCh37), and ClinVar release 2025-10-06 underpin the present version. This per-assembly version split for gnomAD reflects the upstream data landscape: gnomAD releases from v3.0 onward (including the current v4.1) are GRCh38-only, so GRCh37 builds of MetaDome remain anchored to gnomAD r2.0.2 for population variation. ClinVar publishes assembly-specific releases, and the 2025-10-06 release is used for both GRCh37 and GRCh38. UniProtKB/Swiss-Prot and Pfam operate on protein sequence and are therefore assembly-independent, whereas GENCODE, like gnomAD, is tied to a specific assembly.

#### Mapping pipeline

MetaDome maintains a per-nucleotide mapping that links chromosomal positions, through transcript codons, to amino-acid positions in canonical and isoform Swiss-Prot protein sequences and to the Pfam domains containing them, as described previously in MetaDome 2019 release (8). For the 2027 version the mapping is regenerated by a standalone pipeline implemented in Python and orchestrated through a Pixi (22) environment that pins every tool and dependency, and it is run independently for each assembly. GENCODE coding sequences are re-translated and required to reproduce the distributed protein sequence, with in-frame length and a terminal stop codon enforced. Protein-to-protein assignment uses BLAST+ v2.16.0 (23), retaining only pairs that are identical over the full length of both the GENCODE translation and the Swiss-Prot sequence, a stricter criterion than identifier-based cross-references, and stricter than local sequence identity alone. Pfam domains are assigned with pfam_scan v1.6 on HMMER 3.4 (24), applying Pfam gathering thresholds and clan competition, and InterPro annotations are inherited from UniProt’s per-protein cross-references. Per-residue genomic positions are recorded on the transcript’s own strand, separately for each assembly. The pipeline is published at github.com/laurensvdwiel/metadome.

Meta-domains are sets of Pfam domain instances in the human proteome sharing a family annotation, across which variation can be aggregated when a family has at least two human instances. The Swiss-Prot sequences of each family are aligned against its Pfam HMM, producing a Stockholm alignment whose consensus columns define equivalent positions across paralogues. Consensus positions are stored for every family annotated in the human proteome, and aggregation applies to those with two or more instances. In 2019 this aggregation was performed at query time, by retrieving the relevant Stockholm file and mapping each aligned column back to its genomic positions on demand. Here the consensus positions and their per-residue genomic linkages are pre-computed at build time and stored as relational entries (**Supplementary Figure S1**), so per-position aggregation of gnomAD and ClinVar variation is served from indexed lookups. The Stockholm alignments are intermediate build artefacts and are not retained, as the tabular resource carries the same positional correspondence.

We split the database construction into two stages to improve reproducibility and to decouple data generation from platform deployment. Stage one generates a standardised tabular file per genome assembly containing every per-residue mapping between transcript, protein, and Pfam domain, together with the corresponding genomic positions; this file is the canonical mapping resource and is usable independently of the MetaDome platform. Stage two is a batch loader that ingests it into the production PostgreSQL database. Separating these stages allows the underlying mappings to be regenerated, audited, and version-controlled independently of the platform, and allows the platform to be reloaded from an existing tabular resource without re-running the underlying alignments. Both these stage outputs are deposited on Zenodo (25) under CC-BY 4.0, alongside derived per-residue .bed tracks carrying tolerance scores and meta-domain aggregated variation.

Where several Swiss-Prot accessions match the same GENCODE translation at full-length identity, most often paralogous gene-family members with identical canonical sequences, the pipeline retains every candidate and the assignment is resolved at batch loading. The loader prefers the accession whose UniProt gene-name field (GN=) matches the GENCODE gene symbol; where that is not decisive it selects the first-occurring accession deterministically and logs the ambiguity for curation. GN= is used as an auxiliary concordance signal rather than a global exclusion criterion, since GENCODE and UniProt symbols may differ for legitimate reasons such as aliases and paralogous families. The loader enforces a one-to-one transcript-to-UniProt invariant in the production schema (several transcripts may map to one UniProt entry, but each transcript maps to exactly one) and runs pre- and post-load consistency checks for transcript and UniProt key completeness, canonical assignment per transcript, and gene-protein linkage; unresolved cases are reported explicitly rather than silently merged (**Supplementary Table S1**). Row accounting is exact in both builds: every input row is either loaded or recorded as skipped, and all integrity checks returned zero violations.

Competing Swiss-Prot accessions are rare. A single transcript key in 45,778 presented multiple candidates in the GRCh38.p14/GENCODE v45 build, and seventeen in 43,143 in GRCh37.p13/GENCODE v19; in every case the competing accessions were paralogues of identical sequence, which gene-name concordance cannot separate, and the deterministic tie-break was applied (**Supplementary Table S2**). The non-canonical nucleotide-level rows skipped during loading, 1,107 in GRCh38 and 18,057 in GRCh37, are accounted for entirely by these eighteen transcripts, the difference between builds reflecting the number of ambiguous loci rather than any systematic property of the older annotation. Gene-name disagreements between GENCODE and UniProt were retained but flagged for audit rather than excluded, affecting 563 transcripts (1.23%) in GRCh38 and 3,161 (7.33%) in GRCh37 (**Supplementary Table S3**).

The higher mismatch rate in GRCh37 reflects gene-symbol drift between GENCODE v19, released in 2013, and current UniProt nomenclature at release 2025_01. Of 17,928 ENSG identifiers shared between v19 and v45, 972 carry different gene-name labels in the two GENCODE releases (**Supplementary Table S3b**), and a subset of these propagates into GENCODE-versus-UniProt GN= disagreement once symbols were updated in UniProt.

#### Database growth

The MetaDome mapping database integrates the most recent releases of all upstream resources. Across the GRCh38 build, 19,209 protein-coding genes (compared with 18,736 in MetaDome 2019) and 45,778 protein-coding transcripts (42,116) are mapped to 33,629 Swiss-Prot canonical or isoform sequences (33,492) at 100% identity (**Table 1**). On the directly comparable GRCh37 build, annotated Pfam protein-domain instances grew from 71,419 to 82,069 (a 14.9% increase), and Pfam families with at least two within-human occurrences, the families suitable for meta-domain construction, from 3,334 to 5,778 (a 73.3% increase). The GRCh38 build contributes 81,977 instances across 5,706 such families. Pfam annotations are recorded per protein and proteins are shared between builds, so the two builds together hold 87,814 instances across 5,908 families rather than the sum of their parts. Pfam annotations now cover 33,184 of 35,934 (92%) proteins in the resource. At the nucleotide level, 15,291,611 distinct GRCh38 positions (45.7%) and 14,713,557 distinct GRCh37 positions (45.8%) have a Pfam consensus position. Restricting to families with at least two human instances, across which variation can be aggregated, gives 14,302,904 GRCh38 positions (42.7%) and 13,762,784 GRCh37 positions (42.9%). After re-mapping ClinVar release 2025-10-06 onto the updated meta-domain definitions, 67.9% (37,692 of 55,548) of Pathogenic or Likely Pathogenic missense variants in GRCh38 fall at a position within a meta-domain. The equivalent analysis on GRCh37 gives 65.4% (36,316 of 55,542), against 72% (34,076 of ∼47,300) reported for MetaDome 2019 on ClinVar release 2018-05-03. On GRCh37, the number of covered variants increased 6.6%, while ClinVar’s Pathogenic and Likely Pathogenic missense content grew by ∼17% (**Supplementary Table S4**).

**Table 1.** Database content of MetaDome 2027 versus MetaDome 2019 (GRCh38 and GRCh37 builds). Counts of mapped genes, transcripts, Swiss-Prot proteins, Pfam protein-domain instances, meta-domain-eligible Pfam families, distinct chromosome positions, and mapping-table records. Pfam annotations are keyed on protein and proteins are shared between builds, so the per-build Pfam rows overlap and do not sum to the combined totals of 87,814 instances and 5,908 families. The GRCh37 build maps 197 fewer genes than MetaDome 2019 but 1,027 more transcripts, reflecting Swiss-Prot turnover under an unchanged GENCODE v19.

| Data Point | MetaDome 2027<br>(GRCh38) | MetaDome 2027<br>(GRCh37) | MetaDome 2019<br>(<u>only</u> GRCh37) |
| --- | --- | --- | --- |
| Distinct Gencode Gene IDs | 19,209 | 18,539 | 18,736 |
| Distinct Gencode Transcription IDs | 45,778 | 43,143 | 42,116 |
| Total Proteins | 33,629 | 33,937 | 33,492 |
| Pfam protein-domain instances | 81,977 | 82,069 | 71,419 |
| Meta-domain-eligible Pfam families<br>($\geq 2$ human occurrences) | 5,706 | 5,778 | 3,334 |
| Distinct Chromosome and<br>Chromosome Position | 33,479,911 | 32,115,799 | 32,595,355 |
| Distinct Chromosome, Chromosome<br>Position, and Strand | 33,483,493 | 32,120,303 | 32,601,006 |
| Distinct Chromosome, Chromosome<br>Position, and Protein ID | 57,875,359 | 57,855,085 | 57,822,103 |
| Distinct Chromosome, Chromosome<br>Position, and Gene ID | 74,974,791 | 70,673,601 | 70,261,143 |
| Distinct nucleotides with a<br>meta-domain position | 15,291,611 | 14,713,557 | N/A |
| Distinct nucleotides in a Pfam family<br>with $\geq 2$ instances | 14,302,904 | 13,762,784 | N/A |

#### Tolerance landscape

Per-residue tolerance is quantified as the missense-over-synonymous ratio computed from observed gnomAD counts, corrected by the codon-table background of possible missense and synonymous changes, identical in formulation to the original MetaDome (8) and conceptually equivalent to the Missense Tolerance Ratio (6). The tolerance landscape is computed as a sliding 21-residue window across each protein. For GRCh38, recomputation against gnomAD v4.1, assembled from substantially more sequenced individuals than the r2.0.2 release used in MetaDome 2019, increases the observed variant counts underlying each window and correspondingly reduces the sampling noise in the ratio. The GRCh37 landscape retains gnomAD r2.0.2, as no later release targets that assembly. The redesigned pipeline preserves the published GRCh37 tolerance landscape: 99.88% of codons shared with the previous release carry an identical score (**Supplementary Table S5**). Because the two builds draw on different gnomAD releases, tolerance values for the same residue differ between assemblies at nearly all shared protein positions, with only 2.17% identical to within 1e-9.

### Implementation

The MetaDome platform is implemented in Python 3.13 using the Flask web framework and SQLAlchemy 2.0 for relational data modelling. The architecture continues to follow a domain-driven design with rich data entities cached after first construction to minimise per-request database overhead. The full stack is containerised via Docker Compose: a Flask application container, a PostgreSQL v18 database, a Celery v5 task queue for asynchronous retrieval, a Redis v8 result store, a RabbitMQ v4 broker, and a Postfix mail container that handles automated error email notifications and user-initiated support request emails. All version upgrades, the migration from Python 3.5 to 3.13, and the supporting software-stack matrix are listed in **Supplementary Table S6**. Source code and deployment instructions are available at github.com/laurensvdwiel/metadome.

The visualisation layer was modernised for both responsiveness and appearance. The interface uses the Bulma v1.0.4 CSS framework, and interactive genomic and proteomic landscapes are rendered with jQuery v3.7.1 and D3.js v7.9.0, upgraded from v4.13.0 in MetaDome 2019. Three classes of optimisation were applied: pre-computed domain layouts, widths and colour assignments cached on first render; and 60 fps debouncing of brush and zoom interactions with explicit visual feedback. Each Pfam family is now assigned a distinct colour in the schematic protein view, to aid disambiguation when a protein contains several domain types. For the largest human protein transcript (*TTN*, ∼34,000 aa, ENST00000591111.1), the full dashboard visualisation loads in 10.2 s against 48.6 s in the 2019 architecture (median of four loads, MacBook Pro M1 Max, Chrome, same GRCh37 transcript on both platforms), a fivefold improvement reflecting database-indexed lookups, the D3.js upgrade and the modernised backend stack.

Reference-assembly handling. GRCh37 and GRCh38 are represented as parallel, independently-indexed datasets within the same PostgreSQL instance. Selection between assemblies is handled at the web/database interaction layer, so that all downstream visualisations, REST endpoints, and exports automatically reflect the user’s selected assembly without code duplication. GRCh38 is the default; GRCh37 remains available for compatibility with the substantial body of historical clinical sequencing data still anchored to that assembly.

### MetaDome 2027 Web Platform Interface and Positional Endpoints

The MetaDome platform is accessed through a redesigned web interface at www.metadome.app. The principal entry point remains the per-protein dashboard, on which the user supplies a gene symbol, selects a transcript and reference assembly, and receives a tolerance landscape, a meta-domain landscape, an interactive schematic protein view annotated with Pfam domains and ClinVar variation, and a per-position information panel (**Figure 1B**). The 2027 update introduces several extensions to the interaction model. First, the per-protein dashboard now uses informative, deep-linkable URLs of the form

`/metadome/dashboard/{assembly}/{gene}/{transcript}`

accepting query parameters that pre-select one or more residues (`?selected_positions=X1,X2,…`) and pre-open a per-position information panel (`?positional_information=X`). To directly open a certain per-position view, a shorter path form is also accepted

`/metadome/dashboard/{assembly}/{gene}/{transcript}/p.{position}`

Here the position is validated against the transcript length and the request is redirected to the canonical URL above, which makes the compact form convenient wherever a link must be quoted in running text or typed by hand. To preserve user privacy, the query parameters are explicitly stripped before any URL is passed to the user-analytics layer, so the platform does not record which residues a clinician has looked up at a position-by-position level (see **User Communications and Security**). Previously, sharing required either a screenshot of the visualisation or a description of how to navigate to the relevant view; this deep linkable URL pattern now captures both the gene/transcript context and the residue selection in a citable form. Second, MetaDome exposes a URL-based position-lookup endpoint at

`/metadome/position/{assembly}/{chr}/{pos}`

which resolves a genomic coordinate to a table of all matching transcript-protein contexts at that position, each row reporting the gene symbol, transcript identifier(s) (with MANE Select annotation where applicable), UniProt accession, protein position (with codon and reference amino acid), and any covering Pfam domain instance(s). Each row provides a direct deep link to the corresponding dashboard view at that residue. For example, a query at chr7:39686713 returns the *RALA* transcript ENST00000005257.7 for the GRCh38.p14 assembly (MANE Select; UniProt P11233) with its covering Ras Family domain (PF00071) (**Figure 1A**), linked directly to a dashboard view of position p.Lys16, the variant of interest in the illustrative use case described below. Under the 2019 architecture, users navigated from a gene symbol through transcript selection and dashboard rendering before being able to inspect the position of interest; the position-lookup entry point now streamlines that workflow into a single coordinate-based query. Third, a universal search bar accepting either a gene symbol (e.g. *RALA*) or a genomic coordinate (e.g. chr7:39686713) is available on every interface, automatically directing the user to the appropriate dashboard or position-lookup view based on the input type and the selected genome build. Fourth, the GRCh38 build now includes the Y chromosome. Parallel coverage of GRCh37 and GRCh38 is offered through a single interface, allowing users to switch reference assemblies without leaving the dashboard.

#### User communications and security

No personal data is stored beyond support request contact-form submissions. Server-to-user communication is handled through a containerised Postfix mail service that sends automated acknowledgements and error notifications. Form endpoints are protected by reCAPTCHA v3, a honeypot field, and application-layer rate limiting. Analytics are collected exclusively through Google Analytics 4, which anonymises IP addresses by default. The full privacy policy, including handling of cookies, third-party services, and cross-Atlantic data transfer under the EU–US Data Privacy Framework, is published at www.metadome.app/metadome/privacy.

### Community Use of MetaDome

Since publication of the original MetaDome in 2019, the resource has accumulated 317 citations as of April 2026 (Google Scholar). To characterise how the community uses MetaDome, we reviewed all of these citations in full text, of which 261 constituted independent peer-reviewed use of the resource, and assigned each to one of six usage categories (**Figure 2; Supplementary Data S1**): (i) support for clinical variant interpretation under the ACMG/AMP framework, including the reclassification of variants of uncertain significance; (ii) gene-disease association studies at the variant level; (iii) protein-domain mechanism studies; (iv) methods development or comparison against other variant-effect predictors; (v) integration into computational analysis pipelines; and (vi) reviews and educational use. The largest single category was gene-disease association at the variant level (127 publications, 49%), followed by clinical variant interpretation (90, 34%); counting all applicable categories rather than the primary assignment alone, clinical variant interpretation was the most broadly represented use case, appearing in 201 publications (77%). Across these clinical applications MetaDome’s per-residue tolerance landscape was used to provide supporting evidence for the classification of an individual variant of uncertain significance, with 60 publications (23% of reviewed citations) placing that evidence within an explicit ACMG/AMP criterion (26), most frequently PM1 for a variant falling within a mutational hotspot or critical functional domain (23 publications), followed by PM5 (12) and PS1 (9); 11 publications invoked more than one of these three, so 33 in total invoked at least one.

**Figure 2.**
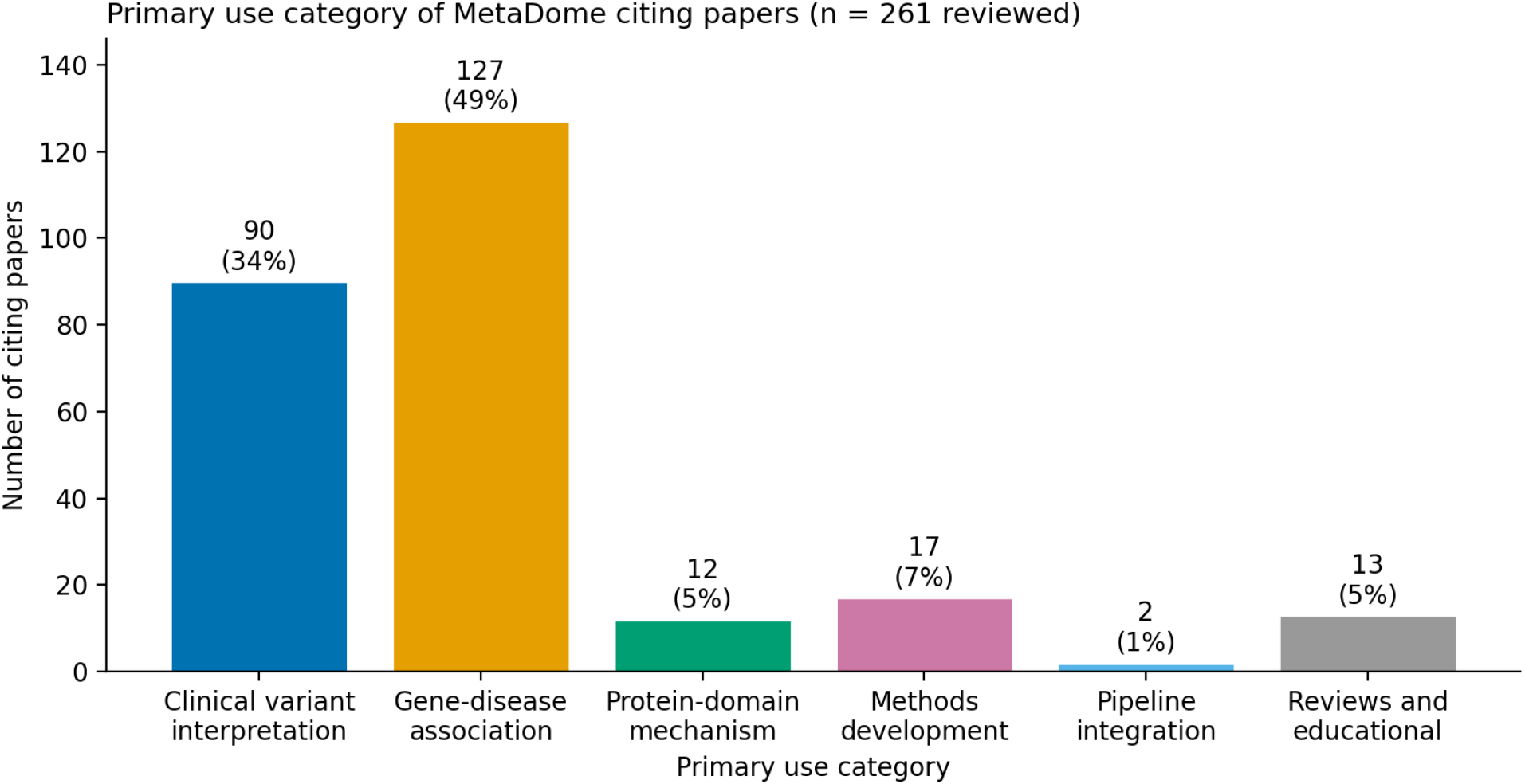
Primary use category of MetaDome 2019 citing papers. Each of the 261 reviewed peer-reviewed publications citing MetaDome 2019 was assigned one primary usage category: clinical variant interpretation (n = 90, 34%), gene-disease association (n = 127, 49%), protein-domain mechanism (n = 12, 5%), methods development or comparison (n = 17, 7%), pipeline integration (n = 2, 1%), and reviews and educational use (n = 13, 5%). Where publications spanned multiple use cases, additional non-exclusive category flags were recorded; per-publication assignments, supporting quotations and full annotation are given in **Supplementary Data S1**.

#### Adoption in clinical variant-interpretation guidelines

MetaDome has been incorporated by name into the operational guidance that national diagnostic genetics communities use to apply the ACMG/AMP framework. The Association for Clinical Genomic Science (ACGS) 2024 Best Practice Guidelines for germline variant classification reference MetaDome under PM1 (27), where a variant in a functional protein domain that lies in a region the MetaDome tolerance landscape calls intolerant supports moderate evidence of pathogenicity, downgradable to supporting where no specific function has been described for the intolerant region. In 2026, the ACGS UK Somatic Variant Interpretation Group (SVIG-UK) guidelines extended this into cancer somatic variant classification, listing MetaDome alongside gnomAD and DECIPHER (28) as recommended sources of regional missense constraint for the new oncogenicity code O8 (29). The ACMG/AMP 2015 sequence-variant interpretation standards themselves (26) predate MetaDome and do not name it directly; the explicit clinical recommendations to consult MetaDome come from ACGS and SVIG-UK rather than from ACMG itself. The Bayesian calibration of paralogue-aggregated evidence by Gunning and Wright (2023) (16) provides quantitative likelihood-ratio support for applying meta-position evidence at moderate weight under PM5, which is consistent with how the ACGS and SVIG-UK guidelines now position MetaDome’s outputs (27, 29). Zhang et al. (2024) (17) further developed Homologous Missense Constraint (HMC), a quasi-amino-acid-resolution constraint score on the meta-domain framework, recommending its application under PP2 (low rate of benign missense in a gene where missense is a common mechanism of disease). Both Gunning and Wright (2023) and Zhang et al. (2024) are static, single-release analyses; MetaDome 2027 provides the live, queryable, visualised counterpart that exposes the underlying paralogue evidence.

### Illustrative Use-Case

To illustrate the MetaDome platform workflow, we describe its application to a single variant of uncertain significance. A *de novo* missense variant in *RALA* (NM_005402.4:c.46A>G; p.Lys16Glu), absent from gnomAD v4.1 and from ClinVar at the time of analysis, was identified through trio genome sequencing of a 12-year-old patient with developmental delay, ptosis, dysmorphic features, congenital cardiac defect, recurrent pancreatitis, and abnormal movements since infancy. The phenotype overlapped Hiatt-Neu-Cooper syndrome (30) [OMIM #619311], for which *de novo* missense variants in *RALA* are an established cause. The variant was subsequently submitted to ClinVar by the reporting laboratory and appears as Likely pathogenic (VCV 2582518) in the 2025-10-06 release used here, which is why it is shown on the annotated codon in **Supplementary Figure S2**. Homologous counts exclude records on the annotated codon itself, so this submission does not contribute to the paralogue evidence below.

Position p.Lys16 lies within the Ras Family domain (Pfam PF00071; **Figure 1B**). The MetaDome per-position panel for this residue shows that it aligns to a Pfam consensus position shared with 124 other human codons in homologous Ras family domain instances (**Supplementary Figure S2**). ClinVar variation at those aligned positions includes six Pathogenic or Likely Pathogenic missense variants at the equivalent residue in two paralogues: in *RIT1*, two Lys>Asn substitutions (VCV 3349333, 581105) and one Lys>Glu (VCV 1695891); and in *KRAS*, two Lys>Asn (VCV 12594, 372705) and one Lys>Gln (VCV 2010214). The *RIT1* Lys>Glu variant is the identical amino-acid substitution to the *RALA* variant. ClinVar additionally records *KRAS* p.Lys5Glu at this position (VCV 12596, eight submissions), also an identical substitution, under the combined Pathogenic/Likely_pathogenic classification; MetaDome admits only discrete Pathogenic and Likely_pathogenic classes, so the evidence shown is a lower bound. Under the Bayesian calibration of Gunning and Wright (2023), the presence of pathogenic missense variation at the equivalent meta-position in a homologous protein corresponds to PM5 applied at moderate weight (LR+ = 7) (16). On the basis of this paralogue-equivalent evidence, in combination with protein structural analysis, and absence from population databases, the clinical diagnostic laboratory reclassified the *RALA* p.Lys16Glu variant from VUS to Likely Pathogenic.

The *RALA* case is not exceptional. Taking every single nucleotide variant in ClinVar 2025-10-06 whose CLNSIG is Uncertain_significance and whose molecular consequence includes missense_variant, and intersecting these with the Pfam domain coverage and meta-domain ClinVar tracks of the MetaDome 2027 release, 1,920,156 such variants are recorded on GRCh38, of which 786,926 (41.0%) fall within a Pfam domain and are therefore positioned within a meta-domain (**Figure 3A**). For 147,835 of those (18.8%), at least one pathogenic or likely pathogenic missense variant has been reported at the evolutionarily equivalent consensus position in a homologous domain elsewhere in the genome. Classified by the evidence available (**Figure 3B**), 3,337 (2.3%) also carry pathogenic evidence at the same residue of the same protein, the configuration calibrated at strong weight; 52,463 (35.5%) carry pathogenic evidence at homologous positions only, with no benign variation at those positions to contradict it, the moderate-weight PM5 configuration illustrated by the *RALA* case; and 92,035 (62.3%) carry both pathogenic and benign variation at homologous positions and are therefore conflicting rather than supportive. Across all candidates, 25,407 are supported by three or more variants classified Pathogenic. The equivalent analysis on GRCh37 identifies 140,374 candidates among 765,455 in-domain variants (18.3%), the difference between assemblies reflecting the larger transcript set annotated by GENCODE v45. ClinVar classifications were matched exactly, so records labelled Pathogenic/Likely_pathogenic were not counted as pathogenic; a variant’s own ClinVar records are excluded from its homologous counts by construction of the release; where a variant falls within several domain placements or protein isoforms, the placement carrying the most pathogenic homologues was retained; and identical amino-acid substitutions, which would constitute PS1-type rather than PM5-type evidence, were not distinguished, so the counts reported are a lower bound on the evidence available at each position. Counts are of distinct pathogenic missense single nucleotide variants at the equivalent consensus position, which may arise at a single homologous residue, as in the case above, or be distributed across several homologous domains; the number of distinct contributing genes is not distinguished. No variant is reclassified here. The analysis quantifies where paralogue-equivalent evidence already exists to be weighed alongside the other criteria a diagnostic laboratory applies; it reads only files from the published data release, is reproducible with bcftools and bedtools, and the script is provided in the MetaDome repository at github.com/laurensvdwiel/metadome/tree/master/scripts/variant_analysis, with the full candidate sets for both assemblies deposited as **Supplementary Data S2 and S3**.

**Figure 3.**
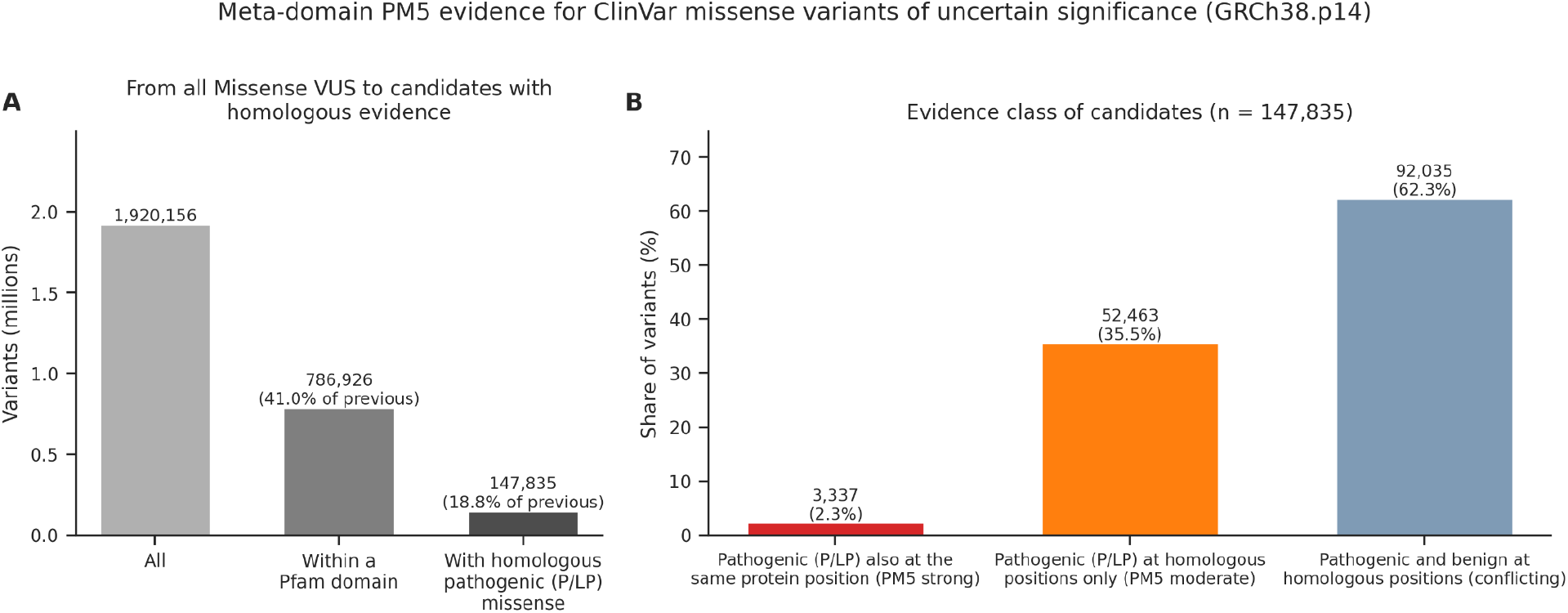
Meta-domain PM5 evidence for ClinVar missense variants of uncertain significance, GRCh38.p14. (A) Of all missense variants of uncertain significance in ClinVar release 2025-10-06 (n = 1,920,156), 786,926 (41.0%) fall within a Pfam domain, and 147,835 of those (18.8%) have at least one pathogenic or likely pathogenic missense variant at the evolutionarily equivalent meta-domain position elsewhere in the genome. (B) The 147,835 candidates by evidence class: 3,337 (2.3%) also have pathogenic evidence at the same residue of the same protein, calibrated at strong weight; 52,463 (35.5%) have pathogenic evidence at homologous positions only, uncontradicted by benign variation at those positions, the moderate-weight PM5 configuration; and 92,035 (62.3%) have both pathogenic and benign variation at homologous positions and are therefore conflicting. Classes are mutually exclusive and each variant is counted once, under the strongest placement available. The equivalent GRCh37 analysis is shown in **Supplementary Figure S3**.

## Discussion

Several directions are anticipated for future MetaDome releases. First, the migration of Pfam to InterPro and the increasing use of AlphaFold-derived structural information for protein-domain boundary refinement (18, 21) will refine MetaDome’s meta-domain definitions and expand the set of recognised human protein domains, partly addressing the limits of purely sequence-based domain prediction. Second, the URL-based position-lookup endpoint is a natural integration point for deep-learning per-variant predictors. Extending it to expose predictions from AlphaMissense (9), EVE (10), and PrimateAI-3D (11) alongside the existing meta-domain context would allow users to consult model-derived predictions and paralogue-aggregated evidence in a single query. Beyond programmatic pipeline integration, the deep-linkable URLs also make MetaDome machine-addressable for AI-based assistance. As variant-curation workflows increasingly incorporate large-language-model agents to draft and cross-check classification reports, a stable, URL-addressable evidence surface allows such agents to point to MetaDome at a specific genomic position for immediate analysis and cite the retrieved result. Third, the Zenodo-deposited mapping resource (25) is designed to support reuse by third parties to incorporate the data underlying MetaDome directly without round-tripping through the platform.

The fundamental approach of MetaDome, aggregating per-position evidence across structurally and evolutionarily related sites within the human proteome, both sits alongside model-based variant scoring as a complementary, traceable evidence source, and increasingly serves as a substrate for refined statistical formulations of the meta-domain signal (16, 17) and for the next generation of model-based predictors. Where model-based methods provide a single per-variant score, MetaDome surfaces the underlying domain homologue evidence in a form that maps directly onto the ACMG/AMP framework, supporting traceable evidence for PM1, PM5, and PS1 reasoning. The 2027 update positions MetaDome to remain a useful resource alongside this rapidly developing predictor landscape.

## Acknowledgements

We thank all research individuals in the UDN and GREGoR consortium. We would also like to thank members of the Wheeler lab, the Montgomery lab, the GREGoR consortium, the UDN, and the Stanford Center for Undiagnosed Diseases, who gave invaluable feedback and assistance throughout this project. We are grateful to Dave Lawrence, Ying Zhu, and Tony Roscioli for engaged use of the platform and constructive feedback over the years. We also thank Maximilian Haeussler, Luis Nassar, and Jairo Navarro of the UCSC Genome Browser team for facilitating the MetaDome track integration into the UCSC Genome Browser. We thank the patient and family described in the illustrative use case for their participation and consent.

AI-assisted coding tools (Claude Code and OpenAI Codex) were used to assist and assess code in the MetaDome repository, which has been in development since 2017; suggested changes and modernisations were incorporated only after manual review by the authors. Large language models (Anthropic Claude) were used to cross-check values and supplementary data derived from the database and the Zenodo deposition for internal consistency, and to assist with wording and grammar. The authors reviewed and verified all AI-assisted outputs and take full responsibility for the content of the manuscript and the software.

## Author Contributions

**Conceptualization:** LW, CG, MTW. **Methodology:** LW, FF. **Software:** LW, FF, JY, JZ, MvdV. **Validation:** LW, RM, CMR, JLC, DEB, JNC, SM, SBM. **Formal analysis:** LW, FF. **Investigation:** LW, FF, CMR, MvdV. **Resources:** JY, JZ, RM, EK, MBN, TWvH, TK, JAB, SBM, CG, MTW. **Data curation:** LW, FF, DN, JLC. **Writing — Original Draft:** LW. **Writing — Review and Editing:** All authors. **Visualisation:** LW. **Supervision:** LW, TWvH, TK, EAA, SBM, MTW. **Project Administration:** LW, MTW. **Funding Acquisition:** LW, FF, TWvH, TK, EAA, JAB, SBM, MTW.

## Data Availability

The MetaDome 2027 web platform is freely available, without registration or login, at www.metadome.app. The source code, the data-construction pipeline, and Docker deployment instructions are available at github.com/laurensvdwiel/metadome under the MIT License. The MetaDome 2027 data release is publicly archived at Zenodo (25) under CC-BY 4.0. For both GRCh37 and GRCh38 it comprises the per-residue transcript-to-protein-to-domain mapping databases including meta-domain consensus linkages, the complete per-position dataset of tolerance scores and aggregated gnomAD and ClinVar variation, the genome-browser tracks derived from that dataset, and the prebuilt output required to reproduce the platform without repeating the build.

