## Supplementary Information for "MetaDome 2027: a comprehensively updated resource for aggregating missense variant evidence across homologous human protein domains"

### MetaDome 2027 Supplementary Information

This document covers Supplementary Data, Tables, and Figures

### Supplementary Data

Supplementary Data is submitted as separate files, here are the descriptions.

**Supplementary Data S1.** Full annotation of the citation review. One row per publication citing MetaDome, covering the 317 citations recorded by Google Scholar as of April 2026, of which 261 were judged independent peer-reviewed uses of the resource and assigned a primary usage category. For each publication the table records the bibliographic details, whether it was included, the primary usage category, any additional non-exclusive category flags where a publication spanned several use cases, the ACMG/AMP criterion invoked where one was named explicitly, and the supporting quotation from the full text on which the assignment was based. Category counts are summarised in **Figure 2**.

**Supplementary Data S2.** ClinVar missense variants of uncertain significance with pathogenic evidence at homologous meta-domain positions, GRCh38.p14. The file contains 368,438 rows covering 147,835 distinct ClinVar variants, because a variant lying in more than one Pfam domain placement or protein isoform appears once per placement; the counts reported in the main text and Figure 3 are of distinct variants, taking for each the placement carrying the most pathogenic homologues. Columns are: chrom, pos, clinvar_id, ref, alt, clnsig and clnrevstat as recorded in ClinVar release 2025-10-06; clnhgvs, the ClinVar genomic HGVS description; gene_symbol, uniprot_ac and uniprot_pos, the protein context of the placement; pfam_id and consensus_pos, the Pfam family and the meta-domain consensus position the residue aligns to; homolog_missense_P_count and homolog_missense_LP_count, the numbers of distinct Pathogenic and Likely_pathogenic missense single nucleotide variants at that consensus position in other genes; clinvar_P_accessions and clinvar_LP_accessions, the corresponding ClinVar variation identifiers; homolog_benign_count, the equivalent count of benign and likely benign variants, which separates supportive from conflicting evidence; same_residue_PLP_count, pathogenic evidence at the same residue of the same protein, the strong-weight configuration in **Figure 3B**; and metadome_url, a direct link to the MetaDome position-lookup view for that coordinate. Records on the annotated codon itself are excluded from the homologous counts by construction of the data release.

**Supplementary Data S3.** As **Supplementary Data S2**, for GRCh37.p13. The file contains 332,525 rows covering 140,374 distinct ClinVar variants. Columns and conventions are identical; genomic coordinates and metadome_url are on GRCh37.p13, and homologous counts are drawn from the GRCh37 build. Summary counts are given in **Supplementary Figure S3**.

### Supplementary Tables

**Supplementary Table S1.** Quality-control metrics for the MetaDome 2027 mapping database, per genome build. Entries created during the GRCh37 import are incremental: GRCh38 was loaded first, so shared proteins, domains and meta-domain positions are created once and reused. Combined totals across both builds: 88,921 gene/transcript entries, 35,934 protein entries, 195,492,981 nucleotide-level mapping rows, 88,702,395 meta-domain mapping entries. Integrity categories checked: genes without protein, mappings without gene, mappings without protein, genes with mappings to multiple proteins, and transcript keys with multiple mapping UniProt accessions.

| QC metric | GRCh38.p14 / GENCODE v45 | GRCh37.p13 / GENCODE v19 |
|---|---|---|
| Input rows | 118,145,565 | 77,366,580 |
| Rows passing all completeness checks | 118,145,565 (100%) | 77,366,580 (100%) |
| Rows with Pfam and meta-domain annotation | 53,251,344 (45.1%) | 35,460,525 (45.8%) |
| Transcript keys | 45,778 | 43,143 |
| Transcript keys with multiple UniProt candidates | 1 | 17 |
| - unresolved by gene-name concordance | 1 | 17 |
| Loaded mapping rows | 118,144,458 | 77,348,523 |
| Skipped non-canonical rows | 1,107 | 18,057 |
| GN= mismatches, transcripts | 563 (1.23%) | 3,161 (7.33%) |
| GN= mismatches, nucleotide rows | 721,869 | 4,659,897 |
| Gene/transcript entries created | 45,778 | 43,143 |
| Protein entries created | 33,629 | 2,305 |
| Pfam/InterPro domain entries created | 81,977 | 5,837 |
| Meta-domain positions created | 1,301,316 | 6,780 |
| Meta-domain mapping entries created | 53,250,759 | 35,451,636 |
| Integrity violations, all categories | 0 | 0 |

**Supplementary Table S2.** All transcript keys with unresolved competing Swiss-Prot accessions, both builds. Skipped rows are the nucleotide-level rows of the non-selected accessions and equal the coding length times the number of skipped accessions; they sum to 1,107 in GRCh38 and 18,057 in GRCh37, the totals reported in Supplementary Table S1. The GRCh38 case and the ENST00000381497 entries in GRCh37 are the same NPIP locus, with the same candidate pair (NPIPA9, A0A0B4J1W7; NPIPA6, P0DXC3) selected identically in both builds. Sequences of competing accessions are identical, so the two NOTCH2NL transcripts and the two RP11-1118M6.1/AC084121.16 transcripts select opposite members of the same candidate pair under the first-occurring tie-break.

| Transcript | Build | GENCODE gene name | Selected UniProt | Skipped UniProt(s) | Skipped rows |
|---|---|---|---|---|---|
| ENST00000381497.6 | GRCh38.p14 | ENSG00000183889 † | A0A0B4J1W7 | P0DXC3 | 1,107 |
| ENST00000362074.6 | GRCh37.p13 | NOTCH2NL | P0DPK4-2 | Q7Z3S9 | 708 |
| ENST00000369340.3 | GRCh37.p13 | NOTCH2NL | Q7Z3S9 | P0DPK4-2 | 708 |
| ENST00000452267.1 | GRCh37.p13 | FAM25B | B3EWG6 | B3EWG3; B3EWG5 | 534 |
| ENST00000381497.2 | GRCh37.p13 | AC138969.4 | A0A0B4J1W7 | P0DXC3 | 1,107 |
| ENST00000331436.4 | GRCh37.p13 | AC138969.4 | A0A0B4J1W7 | P0DXC3 | 1,107 |
| ENST00000427999.2 | GRCh37.p13 | RP11-1212A22.4 | A0A0B4J1W7 | P0DXC3 | 1,107 |
| ENST00000541810.1 | GRCh37.p13 | RP11-1212A22.4 | A0A0B4J1W7 | P0DXC3 | 1,107 |
| ENST00000543226.1 | GRCh37.p13 | TRAPPC2P1 | P0DI82 | P0DI81 | 420 |
| ENST00000596755.1 | GRCh37.p13 | TRAPPC2P1 | P0DI82 | P0DI81 | 420 |
| ENST00000413601.2 | GRCh37.p13 | LIMS3L | P0CW20 | P0CW19 | 351 |
| ENST00000554285.2 | GRCh37.p13 | DUX4 | P0CJ90 | P0CJ89; P0CJ88; P0CJ85; P0CJ86 | 5,088 |
| ENST00000379357.5 | GRCh37.p13 | POLR2J3 | E5RIL1 | B0FP48 | 789 |
| ENST00000340457.8 | GRCh37.p13 | UPK3BL | E5RIL1 | B0FP48 | 789 |
| ENST00000533250.1 | GRCh37.p13 | RP11-1118M6.1 | E9PI22 | P0DMB1 | 1,674 |
| ENST00000528972.1 | GRCh37.p13 | AC084121.16 | P0DMB1 | E9PI22 | 1,674 |
| ENST00000437818.1 | GRCh37.p13 | DEFB130 | P0DP73 | P0DP74 | 237 |
| ENST00000400079.1 | GRCh37.p13 | DEFB130 | P0DP73 | P0DP74 | 237 |

† Recorded as an ENSG identifier in the load report; substitute the GENCODE v45 gene symbol.

**Supplementary Table S3.** Gene-nomenclature concordance. (a) Transcripts whose GENCODE gene symbol disagrees with the UniProt GN= field of the assigned Swiss-Prot entry; retained and flagged, not excluded. (b) Gene overlap between builds; ENSG identifiers are compared with version suffixes stripped.

**(a) Transcript-level GN= mismatch**

| Build | Mismatched transcripts | Total transcripts | Mismatch % |
|---|---|---|---|
| GRCh38.p14 / GENCODE v45 | 563 | 45,778 | 1.23% |
| GRCh37.p13 / GENCODE v19 | 3,161 | 43,143 | 7.33% |

**(b) Gene overlap between builds**

| | Count |
|---|---|
| Gene symbols in both builds | 16,956 |
| Gene symbols only in GRCh38 | 2,229 |
| Gene symbols only in GRCh37 | 1,543 |
| ENSG identifiers in both builds | 17,928 |
| ENSG identifiers in both builds with differing gene symbol | 972 |

**Supplementary Table S4.** ClinVar Pathogenic and Likely Pathogenic missense coverage by meta-domains. Counts are of distinct single nucleotide variants whose ClinVar CLNSIG field is exactly Pathogenic or Likely_pathogenic; the combined Pathogenic/Likely_pathogenic class is excluded. ClinVar publishes assembly-specific releases, so the GRCh38 and GRCh37 totals differ slightly at the same release date. Coverage is the fraction falling at a codon with a populated domain_id and consensus_pos in the deposited dataset, which covers every Pfam family annotated in the human proteome, including families with a single human instance.. The two MetaDome 2027 rows are reproducible from the deposited final dataset and the assembly-matched ClinVar VCF alone, without a database or a running application, by intersecting Pathogenic and Likely Pathogenic missense SNVs with dataset rows whose domain_id and consensus_pos are populated. The MetaDome 2019 total is approximate, recovered from a coverage figure reported to two significant figures in the original publication.

| Build | ClinVar release | P/LP missense SNVs | Within a meta-domain | Coverage |
|---|---|---|---|---|
| GRCh38.p14 | 2025-10-06 | 55,548 | 37,692 | 67.9% |
| GRCh37.p13 | 2025-10-06 | 55,542 | 36,316 | 65.4% |
| GRCh37, MetaDome 2019 | 2018-05-03 | ~47,300 | 34,076 | 72% |

**Supplementary Table S5:** Comparison of the GRCh37 tolerance landscape against the previous MetaDome data release (Zenodo 6625251). Because the GRCh37 build retains gnomAD r2.0.2, the two landscapes are directly comparable. Of the 10,605,455 codons present in both, 99.88% carry an identical tolerance score and the remaining 12,566 differ. The pipeline redesign therefore preserves the published tolerance landscape rather than redefining it. The shared-codon percentage is of the union of codons in the two releases; the identical-score percentage is of the shared codons.

| **What** | **Amount** |
|---|---|
| codons in the 2019 v1.0.1 release | 10,882,508 |
| codons in MetaDome 2027 (GRCh37) | 10,848,533 |
| Shared codons between releases | 10,605,455 (95.3%) |
| Present only in v1.0.1 | 277,053 |
| Present only in MetaDome 2027 | 243,078 |
| **shared codons with identical dn/ds (<1e-9)** | **10,592,889 (99.88%)** |

**Supplementary Table S6**. Software stack of MetaDome 2027 against MetaDome 2019. Versions for 2019 are taken from the repository at commit 484560c (22 May 2019); versions for 2027 are those pinned in the deployed stack and, for the build-time tools, resolved by pixi.lock.

| Layer | Component | MetaDome 2019 | MetaDome 2027 |
|---|---|---|---|
| Runtime | Python | 3.5 | 3.13.3 |
| | Flask | 0.12.4 | 3.1.1 |
| | Werkzeug | 0.14.1 | 3.1.5 |
| | Jinja2 | 2.10.1 | 3.1.6 |
| | gunicorn | 19.8.1 | 23.0.0 |
| Data access | SQLAlchemy | 1.3.0 | 2.0.41 |
| | Flask-SQLAlchemy | 2.3.2 | 3.1.1 |
| | PostgreSQL driver | psycopg2 2.7.3.2 | psycopg 3.3.3 |
| Services | PostgreSQL | 10 | 18.0 |
| | Redis | 4.0 | 8.2.3 |
| | RabbitMQ | 3.7 | 4.2.0 |
| | Postfix mail | not present | boky/postfix 4.4.0 |
| Task queue | Celery | 4.2.0 | 5.5.2 |
| | kombu | 4.2.1 | 5.5.3 |
| | redis client | 2.10.6 | 6.1.0 |
| Scientific Python | NumPy | 1.13.3 | 2.2.5 |
| | pandas | 0.21.0 | 2.2.3 |
| | SciPy | 1.0.0 | 1.15.3 |
| | Biopython | 1.70 | 1.85 |
| | pysam | 0.13 | 0.23.0 |
| | VCF parser | PyVCF 0.6.8 | PyVCF3 1.0.4 |
| | scikit-learn | 0.19.1 | no longer required |

| | | | |
|---|---|---|---|
| Front end | D3.js | 4.13.0 | 7.9.0 |
| | jQuery | 3.3.1 | 3.7.1 |
| | jQuery UI | 1.12.1 | 1.14.1 |
| | Bulma | 0.7.1 | 1.0.4 |
| Build-time tools | BLAST+ | 2.6.0 | 2.16.0 |
| | HMMER | 3.1b2 | 3.4 |
| | pfam_scan | not used | 1.6 |
| | ClustalW | 2.1 | no longer required |
| | Environment management | manual installation | Pixi, with pixi.lock |

### Supplementary Figures

**Supplementary Figure S1.** MetaDome 2027 relational schema. MetaDome is backed by a PostgreSQL database. mappings holds one entry per coding nucleotide, linking a genomic position to its codon and the residue it encodes. genes carries GENCODE transcript annotation, proteins the corresponding UniProtKB/Swiss-Prot entries, and interpro_domains the protein-domain annotation, restricted here to Pfam. New in this release, meta_domain_positions and meta_domain_mapping store the pre-computed correspondence between each Pfam consensus position and every human residue aligned to it, so meta-domain aggregation is served by indexed lookup rather than by parsing alignment files at query time. A genome_build column on genes allows GRCh37 and GRCh38 to coexist in one database, and ext_db_version records the Pfam release each domain annotation came from.

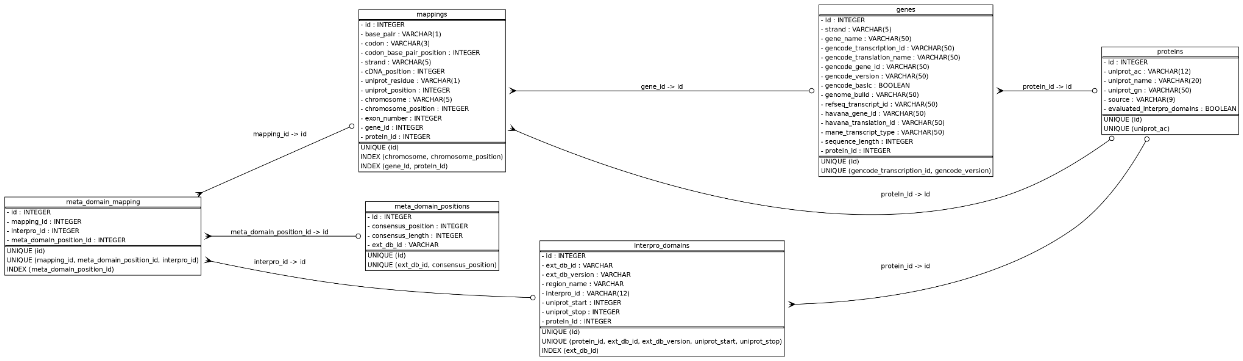

**Supplementary Figure S2.** The MetaDome per-position information panel for RALA p.Lys16. The panel reports the transcript, RefSeq and UniProt identifiers; the residue's genomic interval, codon and exons; its tolerance score (dn/ds 0.37, intolerant) and covering Pfam domain (PF00071, Ras family); ClinVar records on the annotated codon itself; the Pfam consensus position and the 124 other human codons aligned to it; and the Pathogenic and Likely Pathogenic ClinVar records at those homologous codons, here six variants across *RIT1* and *KRAS*. Homologous records exclude the annotated codon, which is listed separately above.
See: metadome.app/metadome/dashboard/GRCh38.p14/RALA/ENST00000005257.7/p.16

### Positional information (p.16)

**Protein details**

Protein of RALA (GENCODE: ENST00000005257.7, RefSeq: NM_005402.4, UniProt: P11233)

**Location details**

Chr: chr7, strand: +
Gene: g.39686713-39686715
Protein: p.16 Lys
cDNA: c.46-48 AAA
Exon: 2, 2, 2
Tolerance score (dn/ds): 0.37 (intolerant)
Position is part of protein domain(s): PF00071

**Known pathogenic ClinVar SNVs at position**

| Gene | Position | Variant | Residue change | Type | Class | ClinVar ID |
|---|---|---|---|---|---|---|
| RALA | chr7:39686713 | A>G | Lys>Glu | missense | LP | 2582518 |

**Meta-domain information for domain PF00071:**

Aligned to consensus position 1, related to 124 other codons throughout the genome (with a 90.6% alignment coverage).

**Pathogenic & Likely Pathogenic ClinVar SNVs at homologous positions:**

| Gene | Position | Variant | Residue change | Type | Class | ClinVar ID |
|---|---|---|---|---|---|---|
| RIT1 | chr1:155910693 | T>A | Lys>Asn | missense | P | 3349333 |
| RIT1 | chr1:155910693 | T>G | Lys>Asn | missense | P | 581105 |
| RIT1 | chr1:155910695 | T>C | Lys>Glu | missense | P | 1695891 |
| KRAS | chr12:25245370 | T>A | Lys>Asn | missense | P | 12594 |
| KRAS | chr12:25245370 | T>G | Lys>Asn | missense | P | 372705 |
| KRAS | chr12:25245372 | T>G | Lys>Gln | missense | LP | 2010214 |

**Supplementary Figure S3**. Meta-domain PM5 evidence for ClinVar missense variants of uncertain significance, GRCh37.p13. Panels as in Figure 3. (A) Of all missense variants of uncertain significance in ClinVar release 2025-10-06 (n = 1,920,209), 765,455 (39.9%) fall within a Pfam domain, and 140,374 of those (18.3%) have at least one pathogenic or likely pathogenic missense variant at the evolutionarily equivalent meta-domain position elsewhere in the genome. (B) The 140,374 candidates by evidence class: 3,179 (2.3%) also have pathogenic evidence at the same residue of the same protein, calibrated at strong weight; 51,479 (36.7%) have pathogenic evidence at homologous positions only, uncontradicted by benign variation at those positions, the moderate-weight PM5 configuration; and 85,716 (61.1%) have both pathogenic and benign variation at homologous positions and are therefore conflicting. Classes are mutually exclusive and each variant is counted once, under the strongest placement available.

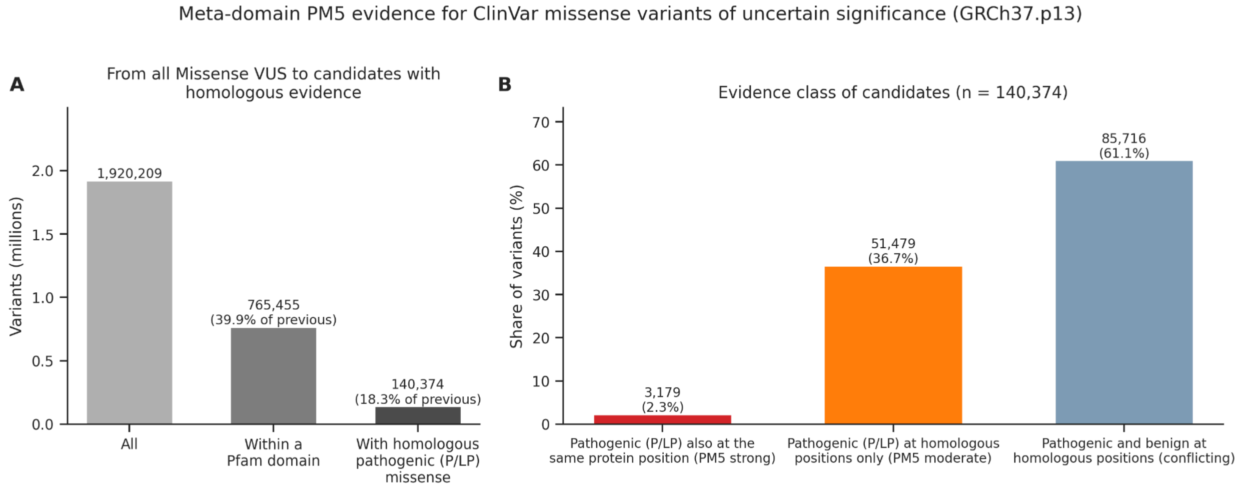
